# Spectrin-based Membrane Mechanics Is Asymmetric and Remodels during Neural Development

**DOI:** 10.1101/2020.04.29.067975

**Authors:** Ru Jia, Yongping Chai, Chao Xie, Gai Liu, Zhiwen Zhu, Kaiyao Huang, Wei Li, Guangshuo Ou

**Affiliations:** Tsinghua-Peking Center for Life Sciences, Beijing Frontier Research Center for Biological Structure, McGovern Institute for Brain Research, School of Life Sciences and MOE Key Laboratory for Protein Science, Tsinghua University, Beijing, China; Key Laboratory of Algal Biology, Institute of Hydrobiology, Chinese Academy of Sciences, Wuhan, China; School of Medicine, Tsinghua University, Beijing, China

**Keywords:** Neural development, Cell migration, Dendrite formation, Cytoskeleton, Membrane mechanics, Spectrin

## Abstract

Perturbation of spectrin-based membrane mechanics causes hereditary elliptocytosis and spinocerebellar ataxia, but the underlying cellular basis of pathogenesis remains unclear. Here, we introduced the conserved disease-associated spectrin mutations into the *C. elegans* genome and studied the contribution of spectrin to neuronal migration and dendrite formation in developing larvae. The loss of spectrin generates an ectopic actin polymerization outside of the existing front and secondary membrane protrusions, leading to defective neuronal positioning and dendrite morphology in adult animals. Spectrin accumulates in the lateral and the rear of migrating neuroblasts and redistributes from the soma into the newly formed dendrites, indicating that the spectrin-based membrane skeleton is asymmetric and remodels to regulate actin assembly and cell shape during development. We affinity-purified spectrin from *C. elegans* and showed that its binding partner ankyrin functions with spectrin. Asymmetry and remodeling of membrane skeleton may enable spatiotemporal modulation of membrane mechanics for distinct developmental events.

**Significance Statement:** The biomechanical regulation of neural development is largely unknown. The spectrin-based membrane skeleton is essential for the structural integrity of the plasma membrane. This study addresses the function and behavior of spectrin in neuroblast migration and dendrite formation. The loss of spectrin generates an ectopic actin polymerization outside of the existing front, leading to defective neuronal positioning and dendrite morphology. Spectrin is absent from the leading edge but accumulates in the posterior of migrating neuroblasts and redistributes from the soma into the nascent dendrites, indicating that the membrane skeleton is asymmetric and remodels. Asymmetry and remodeling of the membrane skeleton may enable spatiotemporal modulation of membrane mechanics for distinct developmental events.

## Introduction

The development of the nervous system requires the correct positioning of neurons and the accurate sprouting and elongation of neurites, laying the foundation for functional neuronal connectivity. The motility of neuronal cell and the growth cone is initiated by the environmental cue-activated transmembrane receptors and followed by asymmetric actin polymerization in the leading edge and the extension of the front of the cell (1-3). Despite the wealth of information on the biochemical mechanism of neural development, ranging from the guidance cues, receptors, the protein machinery that powers motility, the physical regulation of neural development, in particular, the underlying membrane mechanics, is mostly unexplored.

The spectrin-based membrane skeleton is essential for the structural integrity of the plasma membrane and protects cells from mechanical stresses in erythrocyte and many non-erythrocyte cells (4). The functional unit of spectrin is composed of a rod-shaped tetramer that consists of two antiparallel heterodimers of α-spectrin and β-spectrin (4). Spectrin interacts with the actin cytoskeleton and associated proteins to form a static polygonal lattice structure underneath the plasma membrane. Recent super-resolution imaging studies uncovered that spectrin creates an ordered periodic longitudinal array around the circumference of axons and dendrites (3, 5-8). In addition to mechanic support, the membrane-associated periodic skeleton serves as a dynamic signaling platform for G protein-coupled receptors and cell adhesion molecules-mediated receptor tyrosine kinases (9). Mutations in human spectrin genes impair spectrin-based membrane mechanics and cause various diseases. For example, the hereditary elliptocytosis (HE)-associated α-spectrin L260P mutation locks spectrin in the closed dimer conformation and destabilizes the plasma membrane (10, 11). The spinocerebellar ataxia type 5 (SCA5)-associated β-III– spectrin deletion disrupts the spectrin tetramer conformation and alters the localization of the synaptosomal proteins in the cerebellum (12, 13). Knockout of β-II-spectrin in mouse neural progenitors disrupted axonal growth, stability, and synaptic cargo transport (14). These studies establish the importance of spectrin in the nervous system and reveal the association of spectrin deficiency in neurodegenerative disorders; however, the function of the spectrin-based membrane mechanics in neural development remains unclear.

To model spectrin-associated diseases in animals, we have recently introduced the conserved disease-associated mutations into the *C. elegans* α- and β-spectrin genes (15). These mutations affected membrane mechanics, and caused uncoordinated animal movement, mimicking the syndromes of the patients. Using these animals, we uncovered a novel function of spectrin in cilium formation in nematodes and showed the conserved mechanism in mammals (15). Here, we chose the *C. elegans* Q neuroblasts as a tractable experimental system to study the function and behavior of spectrin in developing neural progenitors. The Q neuroblasts, QR on the right side of the animal and QL on the left, undergo an identical pattern of asymmetric cell divisions, directional migration, apoptosis, and neurite growth in the first larval stage (16, 17). The Q cell descendants move to highly stereotypic positions along the anterior and posterior body axis but in opposite directions, with QR and progenies (abbreviated as QR.x) migrating anteriorly and QL and progenies (QL.x) migrating posteriorly (Fig. 1A-B). We developed fluorescence time-lapse microscopy and reporters to study cell polarity and the actin cytoskeleton in migrating Q cells (16, 18-21). Using the system, we show that spectrin is crucial for neuronal migration and dendrite formation and that the spectrin-based membrane skeleton is asymmetrically assembled and remodels in developing neuroblasts.

**Figure 1.**
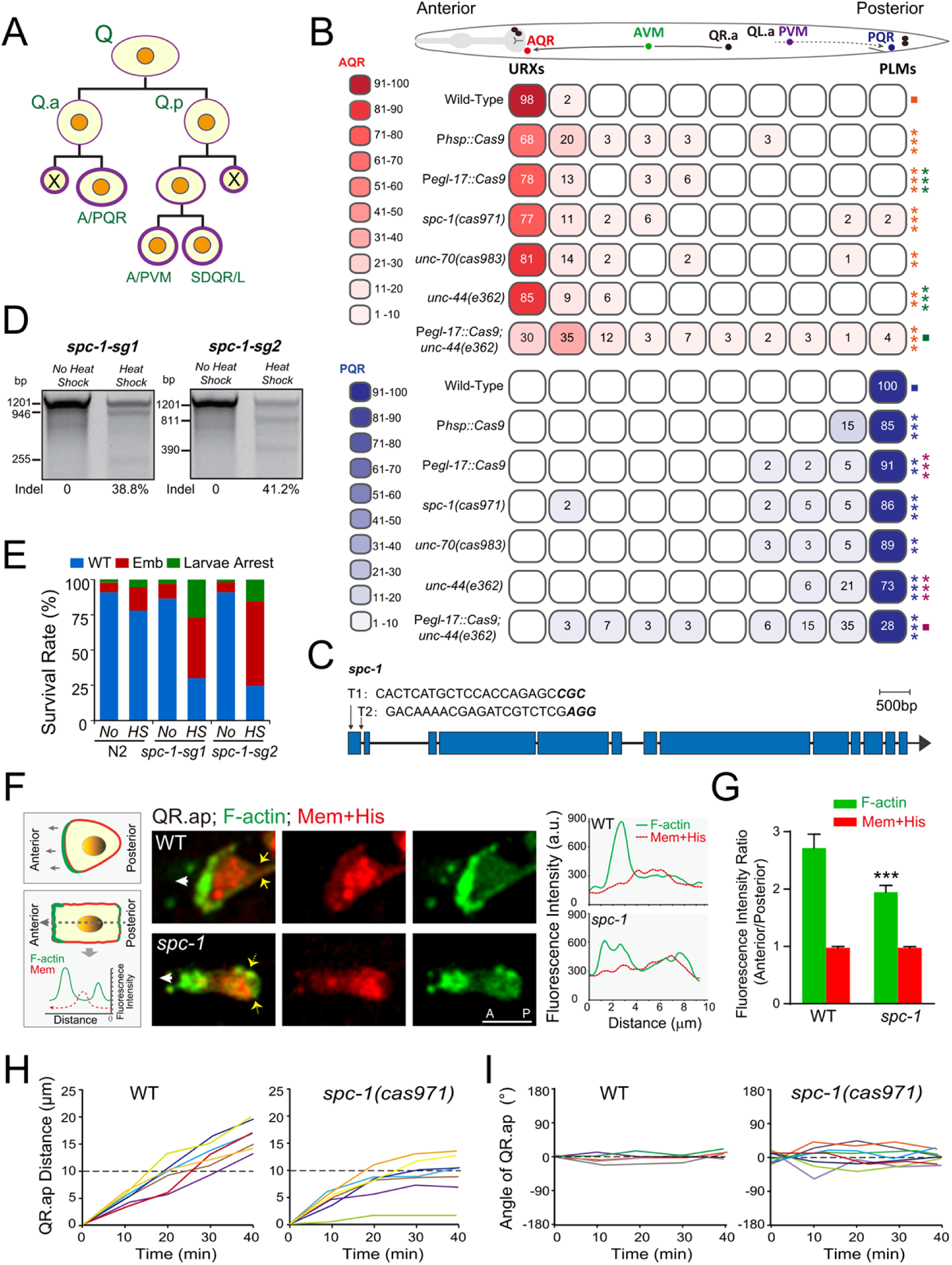
SPC-1 regulates the *C. elegans* Q neuroblast migration. **(A)** Schematic of Q neuroblast lineages. QL and QR neuroblasts divide three times and generate three neurons (QL: PQR, PVM, and SDQL; QR: AQR, AVM, and SDQR) and two apoptotic cells (X). **(B)** A color-coded heatmap is scoring the AQR and PQR position in L4 animals of the indicated genotypes. As indicated in the schematic overview of Q cell migration at the top, the QR descendant (QR.x), AQR, migrates anteriorly, and the QL descendant (QL.x), PQR, migrates posteriorly. The full length between the URXs and the tail is divided into 10 blocks, and the percent of AQRs or PQRs that stopped within each block are listed. The color (red for AQR and blue for PQR) darkness of the blocks symbolizes the range of percentage values. n > 50 for all genotypes. Statistical significance compared to the control (■) with matching color codes is based on χ^2^ test, ** *p* < 0.001, *** *p* < 0.001. **(C)** Schematic of *spc-1* gene model and sgRNA sequences (both sg1 and sg2 target the first exon). **(D)** Representative gels of the T7EI assay for *spc-1* PCR products amplified from the genomic DNA of worms expressing P*hsp::Cas9* and P*U6::spc-1-sg1* (left) or P*U6::spc-1-sg2* (right) with or without heat-shock treatment. **(E)** Quantification of embryo survival rates in WT and *spc-1* conditional knockouts. *N*= 50-100 from three generations. **(F)** Middle: Fluorescence images of GFP-tagged F-actin with mCherry labeled plasma membrane and histone in QR.ap cells in WT or *spc-1(cas971)* animals. Yellow arrows show the rear of migrating cells; White arrows indicate the direction of migration. Anterior, left; Bar, 5 μm. Right: line scans (performed as in the right panel) of fluorescence intensities across the representative migrating cell in the middle panel, respectively. **(G)** F-actin (green) fluorescence intensity ratio of the anterior and to the posterior, the corresponding mCherry-membrane intensity ratio was used as an internal control. The dashed line in Fig. 2A divides the anterior and the posterior. The error bars are the standard error of the mean (SEM). Statistical significance is based on Student’s *t-* test, *** *p* < 0.001. **(H-I)** Quantification of the QR.ap migration distance (**H**), and angle(**I**). Each line in (**H** and **I**) represents the measurement from one time-lapse movie of QR.ap migration.

## Results

### Spectrin regulates neuronal migration by preventing ectopic actin assembly

The germline deletion of α-spectrin (*spc-1)* or β-spectrin (*unc-70*) leads to embryonic lethality (22, 23). To study the function of spectrin in postembryonic neural development, we generated weak loss-of-function alleles or conditional mutations of spectrin. Our early study introduced the disease-associated α-spectrin L260P variation or β-III–spectrin in-frame deletion into the *C. elegans spc-1(cas971)* or *unc-70(cas983)* locus, respectively (15). Both alleles produced viable progenies, allowing the dissection of the role of spectrin in neural development. As a complementary approach, we used the somatic CRISPR-Cas9 method (24) in which the Cas9 endonuclease is expressed under the control of the Q cell-specific promoter P*egl-17* or the heat-shock gene *hsp-16*.*2* promoter P*hsp* to create conditional mutants of *spc-1* within Q cell lineages or during Q cell development. T7 endonuclease I (T7EI)-based assays demonstrated that these transgenic animals produced molecular lesions with the expected sizes at the target loci of *spc-1* after heat-shock induction of Cas9 expression (Fig. 1C-D). In line with the reported phenotypes of the *spc-1(cas971)* animals (15), the *spc-1* conditional mutant embryos but not wild-type (WT) embryos exhibited embryonic lethality and larval arrest (Fig. 1E), demonstrating the success in producing conditional mutations in the *spc-1* locus. In these animals, we examined the final positions of QR.ap and QL.ap, which will become the AQR and PQR neuron after long-distance migration towards the anterior or posterior, respectively. Compared with WT animals, AQR localized at a more posterior position, and PQR confined at a more anterior region in *spc-1(cas971)* or *unc-70(cas983)* or somatic CRISPR-Cas9-created *spc-1* conditional mutants, indicating an essential role of spectrin in cell migration (Fig. 1B and Fig. S1A). The cell migration defects from Q-cell specific *spc-1* mutants suggested a cell-autonomous contribution of spectrin to cell migration.

To understand the cellular mechanism by which the cell migration is reduced in *spc-1* mutants, we performed fluorescence time-lapse analysis of cell morphology and the actin cytoskeleton in migrating Q.ap cells (16). We depicted the plasma membrane and nucleus using mCherry-tagged with a myristoylation signal and a histone and the actin cytoskeleton using GFP-tagged with the actin-binding domain of moesin (GFP::moesin ABD). In WT animals, AQR extends a large lamellipodium and forms a fan-like morphology (Fig. 1F and Fig. S1B). Strikingly, the morphology of the *spc-1*-mutant AQR was perturbed, including the formation of a square-shaped cell without a persistent anterior-posterior polarity during migration (Fig. 1F and Fig. S1B). In WT migrating AQR, GFP::moesin ABD-marked F-actin accumulates in the leading edge but is absent from the lateral and the rear. The *spc-1*-mutant AQR cell contains GFP::moesin ABD fluorescence in the leading edge (i.e., anterior); however, an ectopic GFP fluorescence is detectable in the lateral and the rear (i.e., posterior), resulting in an even distribution of F-actin along the entire plasma membrane (Fig. 1F). The fluorescence intensity ratio between the anterior and the posterior was reduced from 2.7-fold in WT to 1.9-fold in *spc-1* mutant AQR cells (Fig. 1G), indicating a defective organization of the actin cytoskeleton. In agreement with the final reduction of AQR migration in adult animals, AQR failed to persistently extend the lamellipodium and translocate the cell body towards the anterior, reducing the migration distance and persistent polarity over time (Fig. 1H-I). These results show that spectrin is essential for F-actin organization during cell migration. Given the abnormal actin assembly outside of the front, we postulate that the spectrin-based membrane mechanics may act in the posterior to inhibit ectopic actin polymerization.

### Asymmetric distribution of membrane skeleton in migrating neuroblasts

To visualize the cellular distribution of the endogenous spectrin proteins during cell migration, we constructed the knock-in (KI) animals to mark SPC-1 and UNC-70 with GFP (15). We genetically crossed the GFP-spectrin knock-in markers into a transgenic animal that expresses mCherry-labeled plasma membrane and histone within Q cell lineages. GFP-tagged SPC-1 or UNC-70 was absent from the leading edge but accumulated in the posterior of the migrating Q.a, Q.p, Q.ap, and Q.pa cells (Fig. 2A-E and Fig. S1C). Quantification of the fluorescence intensity ratio between the front and the posterior (divided by dashed lines in Fig. 2C-D) showed that GFP-spectrin in the posterior was 2.7 ∼ 2.9-fold brighter than that in the anterior (Fig. 2A-E and Fig. S1C). In both the QL and QR lineages, GFP::SPC-1 or UNC-70::GFP shows a similar enrichment in the posterior of migrating cells (Fig. 2C-D and Movie S1-2). We did not detect any change in red fluorescence of the mCherry-tagged plasma membrane and histone throughout Q cell development (Fig. 2E and Fig. S1C). The spectrin evenly distributes in Q neuroblasts and Q.paa and Q.pap cells that do not undergo long-distance migration (Fig. 2E and Fig. S1C).

**Figure 2.**
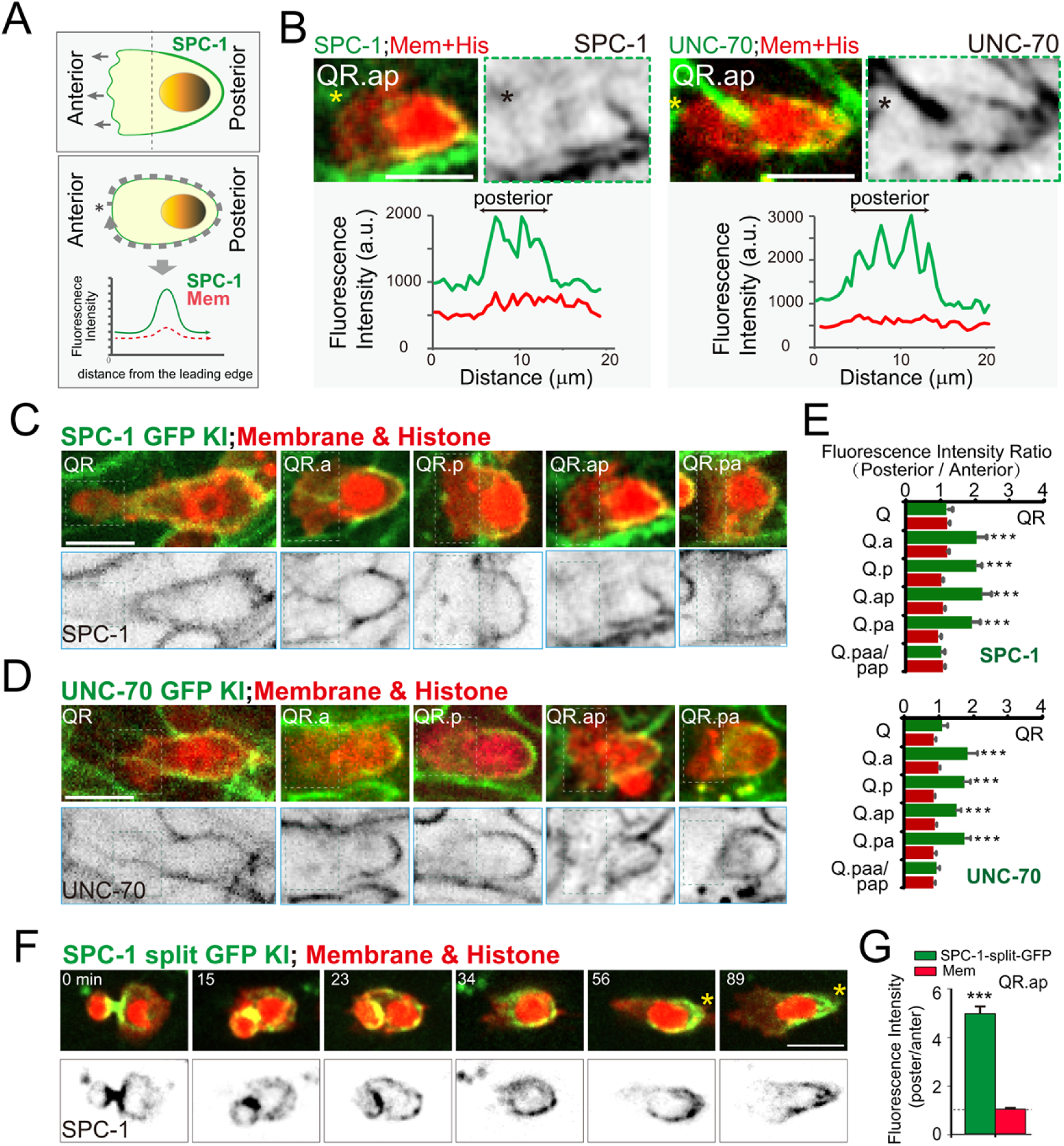
SPC-1 and UNC-70 asymmetrically localize in the migrating cells. **(A)** Upper: schematic of a migrating Q.x cell in SPC-1 GFP-knock-in *C. elegans*, showing GFP fluorescence enrichment at the rear portion of the migrating cell; lower: the outline of GFP fluorescence intensity plot along the migrating cell periphery, starting from the leading spot indicated with asterisk and then along the dashed line around the Q.x cell sketch. **(B)** The representative images of GFP-tagged SPC-1 (left) and UNC-70 (right) in QR.ap cells from knock-in animals. In each panel, upper left showing GFP KI fluorescence (green) with mCherry tagged plasma membrane and histone (red), upper right showing inverted fluorescence images of GFP KI fluorescence, and bottom showing the GFP (green) and mCherry (red) fluorescence distribution plot along the cell periphery of the above representative image. Asterisk, the plot start point. **(C-D)** Fluorescence images of GFP-tagged (**C**) SPC-1 (green) or (**D**) UNC-70 (green) and mCherry tagged plasma membrane and histone(red) in the QR lineage in knock-in animals. lower, inverted images of GFP-knock-in signal of (**C**) SPC-1 and (**D**) UNC-70. **(E)** Quantification of the GFP (green) of SPC-1 (upper) orUNC-70 (lower), and mCherry (red) fluorescence intensity ratio between the leading edge (indicated with the dashed box in **C**-**D**) and the posterior plasma membrane potions in the QR cell lineages (N = 10-20 animals). See Figure S1 for the QL lineage. **(F)** Fluorescence time-lapse images of split-7xGFP-tagged SPC-1, and mCherry-tagged plasma membrane and histone after QR.ap birth in WT. Upper: merged images; lower: inverted 7x-GFP images of SPC-1. **(G)** Quantification of the fluorescence intensity ratio of 7xGFP-tagged SPC-1 (green), and mCherry-membrane (red) between the posterior and the anterior of the migrating QR.ap cells (N = 14 animals). The error bars indicate the standard error of the mean (SEM). Statistical significance is based on Student’s *t-*test, *** *P* < 0 .001. Scale bar in B, C, D, and F, 5 μm.

We sought to pinpoint how spectrin became asymmetrically localized in migrating Q cells. The GFP-spectrin knock-in strains illuminate spectrin in Q cells and their neighboring tissues, the high background fluorescence of which impedes the visualization of spectrin dynamics in Q cells. To resolve this problem, we adopted a self-complementing split fluorescent protein system (15, 25). We inserted seven copies of GFP11 tag that encoded the eleventh beta-strand of super-folder GFP into the *spc-1* locus and expressed GFP1-10 under the control of a Q cell-specific promoter P*egl-17*. After genetically crossing an SPC-1::7XGFP11 knock-in animal with a P*egl-17*::GFP1-10 transgenic animal, GFP self-complemented and specifically illuminated spectrin with green fluorescence in Q cells. Using the strain, we confirmed that SPC-1::7XGFP fluorescence asymmetrically localized in the lateral and the rear of the migrating Q.ap cells, the enrichment fold of which elevated from 2.7-fold in GFP knock-in animals to 5.2-fold in Q-cell specific 7XGFP knock-in animals (Fig. 2G), possibly because of an increase of green fluorescence and elimination of background signal. Next, we followed the dynamic changes of spectrin during Q cell development. After Q cell division, spectrin accumulates at the cleavage furrow between two daughter cells (0 min, Fig. 2F, Movie S1). Spectrin then forms puncta outside of the front (23 min) and becomes enriched in the posterior (34 – 89 min), indicating an asymmetric assembly of the spectrin-based membrane skeleton in migrating cells (Fig. 2G and Fig. S1D).

### Membrane skeleton remodels to safeguard dendrite formation

Our fluorescence time-lapse microscopy revealed that the timing and position of sprouting dendrites couples with cell migration. After the A/PQR neuron reaches its destination, the cell body stops, but the same leading-edge continues to elongate, transforming into the growth cone that forms the dendrite (Movie S2) (19, 26). We examined whether and how the spectrin-based membrane mechanics regulates dendrite growth. In 10∼30% of the *spc-1(cas971)* or *unc-70(cas983)* mutant animals, the A/PVM and A/PQR neurons developed neurites whose positioning and extent of growth were perturbed, and the neurites were defective in the formation, guidance or outgrowth (Fig. 3A-C). In WT animals, A/PVM neurons first extend their neurites in the dorsal/ventral (D/V) direction and then elongate along the anterior/posterior (A/P) body axis, and the PQR dendrite sprouts toward the posterior, with the axon growing toward the anterior. In the spectrin-deficient neurons, AVM improperly extended its axon to the posterior, the PVM axon skipped the initial D/V growth but underwent abnormal A/P growth, and PQR split its dendrite (red arrows in Fig. 3B).

**Figure 3.**
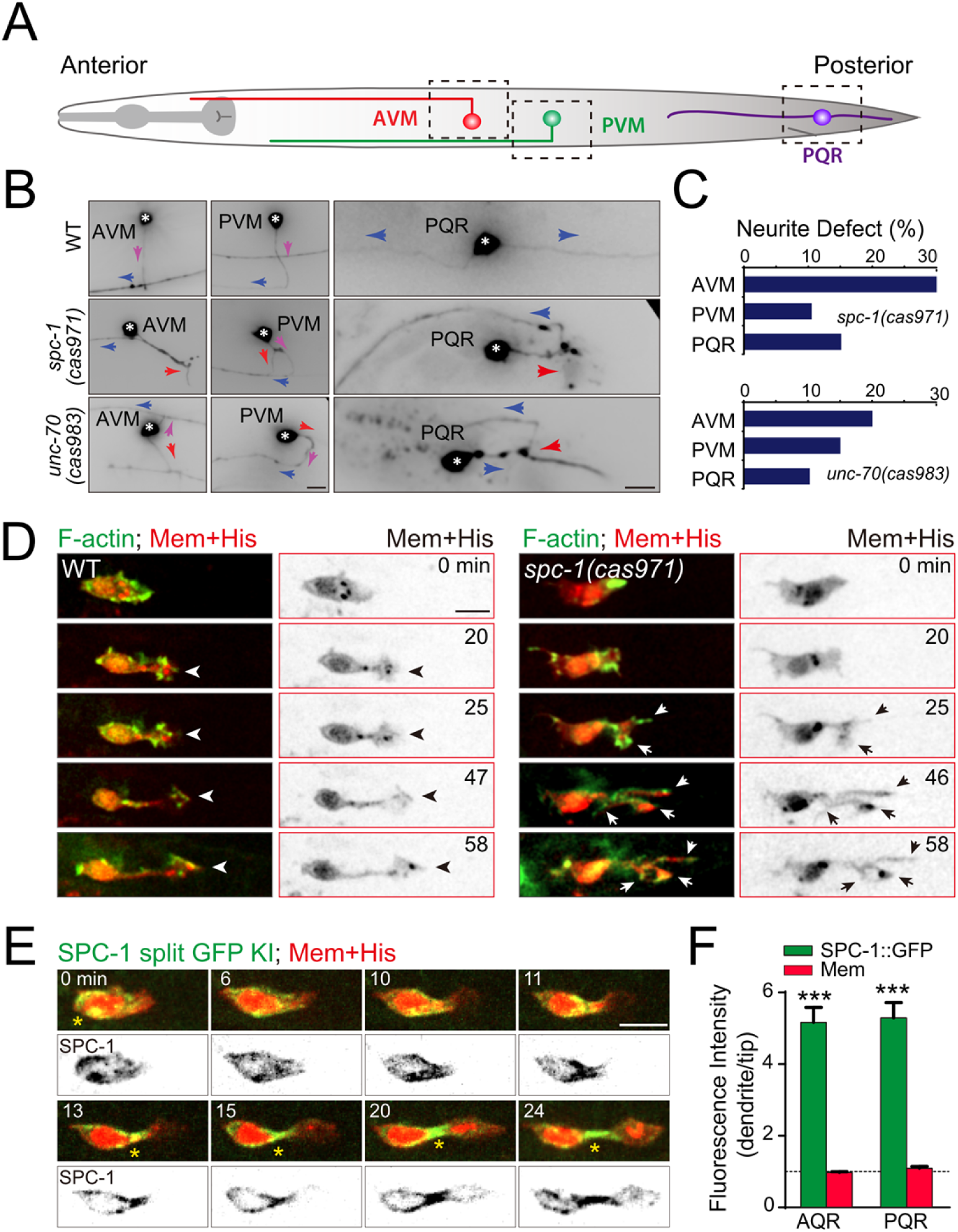
SPC-1 regulates dendrite formation. **(A)** Schematic of the axon and dendrite morphologies of Q cell derived neurons AVM, PVM, and PQR in WT animals. The images of regions indicated by the dashed box are shown in **(B). (B)** Neurites from AVM, PVM, and PQR neurons in WT (upper), *spc-1(cas971)* (middle), and *unc-70(cas983)* (lower) mutant animals. The pink or blue arrows indicate the dorsal-ventral or anterior-posterior growth of the neurites in WT animals, and the red arrows show the misoriented neurites. Asterisk, cell body. **(C)** Quantification of defects in Q cell neuritogenesis in *spc-1(cas971)* (upper) and *unc-70(cas983)* (lower) mutant animals (*N* = 60-100). **(D)** Fluorescence time-lapse images of GFP-tagged F-actin and mCherry-tagged plasma membrane during QL.ap dendrite formation in WT and *spc-1(cas971)* mutant animals, respectively. In each panel, left: merged images; right: inverted mCherry images of cell morphology. asterisk, nucleus; arrow-head, the normal dendrite outgrowth direction towards the animal tail; arrow, the misoriented dendrite in *spc-1(cas971)* mutant animals. **(E)** Fluorescence time-lapse images of 7xGFP-tagged SPC-1, and mCherry-tagged plasma membrane and histone during AQR neuritogenesis in WT. Upper: merged images; lower: inverted split-7x-GFP images of SPC-1. The yellow asterisk, the enrichment of 7xGFP-tagged SPC-1. **(F)** Quantification of the fluorescence intensity ratio of 7xGFP-tagged SPC-1 (green), and mCherry-membrane (red) between the new dendrite and growth cone of the AQR or PQR cells (*N* = 10-14 animals). The error bars indicate the standard error of the mean (SEM). Statistical significance is based on Student’s *t-*test, *** *P* < 0 .001. Scale bar in B, 50 μm; scale bar in D and E, 5 μm.

We performed live imaging analysis of dendrite formation in PQR to address how spectrin regulates dendrite development. In WT animals, the PQR cell body arrived at the final destination, and the leading-edge continued to elongate, forming the growth cone for dendrite formation (Fig. 3D and Movie S2). The F-actin accumulated at the leading edge, and the newly elongated dendrite became thin and stable (Fig. 3D). In spectrin-mutants, the nascent dendrite was unstable and developed the F-actin-enriched secondary growth cones that elongated and generated ectopic dendritic branches (Fig. 3D and Movie S3).

To understand spectrin behavior during dendrite formation, we followed SPC-1::7XGFP in the growing dendrite of the WT PQR neuron: SPC-1 initially accumulated in the lateral and the rear of the migrating PQR; however, SPC-1 transited from the soma to the newly formed dendrite and became enriched in the dendrite (Fig. 3E). We did not detect SPC-1::7XGFP in the leading edge of the growth cone during dendrite outgrowth (Fig. 3E-F), which is consistent with the absence of spectrin in the leading edge of migrating cells. Quantification of the SPC-1::7XGFP ratio between the dendrite and the leading edge revealed a 5.3-fold enrichment (Fig. 3F). Together, these results show that the spectrin-based membrane skeleton disassembles in the soma and assembles in the nascent dendrite, where spectrin inhibits ectopic actin polymerization and safeguards dendrite morphology.

### Ankyrin functions with spectrin during neural development

To dissect the molecular mechanism of how the spectrin-based membrane skeleton regulates neuronal cell migration and neurite growth, we used affinity purification and mass spectrometry to identify interacting proteins of SPC-1. Using an anti-green fluorescent protein (GFP) antibody, we purified GFP-tagged SPC-1 with its associated proteins from the lysate of SPC-1::GFP knock-in larvae (Fig. 4A). We determined protein constituents employing liquid chromatography-tandem mass spectrometry (LC-MS) (Fig. 4B). We detected the beta-spectrin subunit UNC-70, validating our purification scheme. Another hit was UNC-44, an ortholog of human ankyrin protein, which was a binding partner of the membrane skeleton (27). In *C. elegans*, UNC-44 regulates multiple neuronal events, including axon guidance, microtubule organization, and axon-dendrite sorting and transport (28, 29), but its functions in neuronal cell migration and dendrite development are less understood.

**Figure 4.**
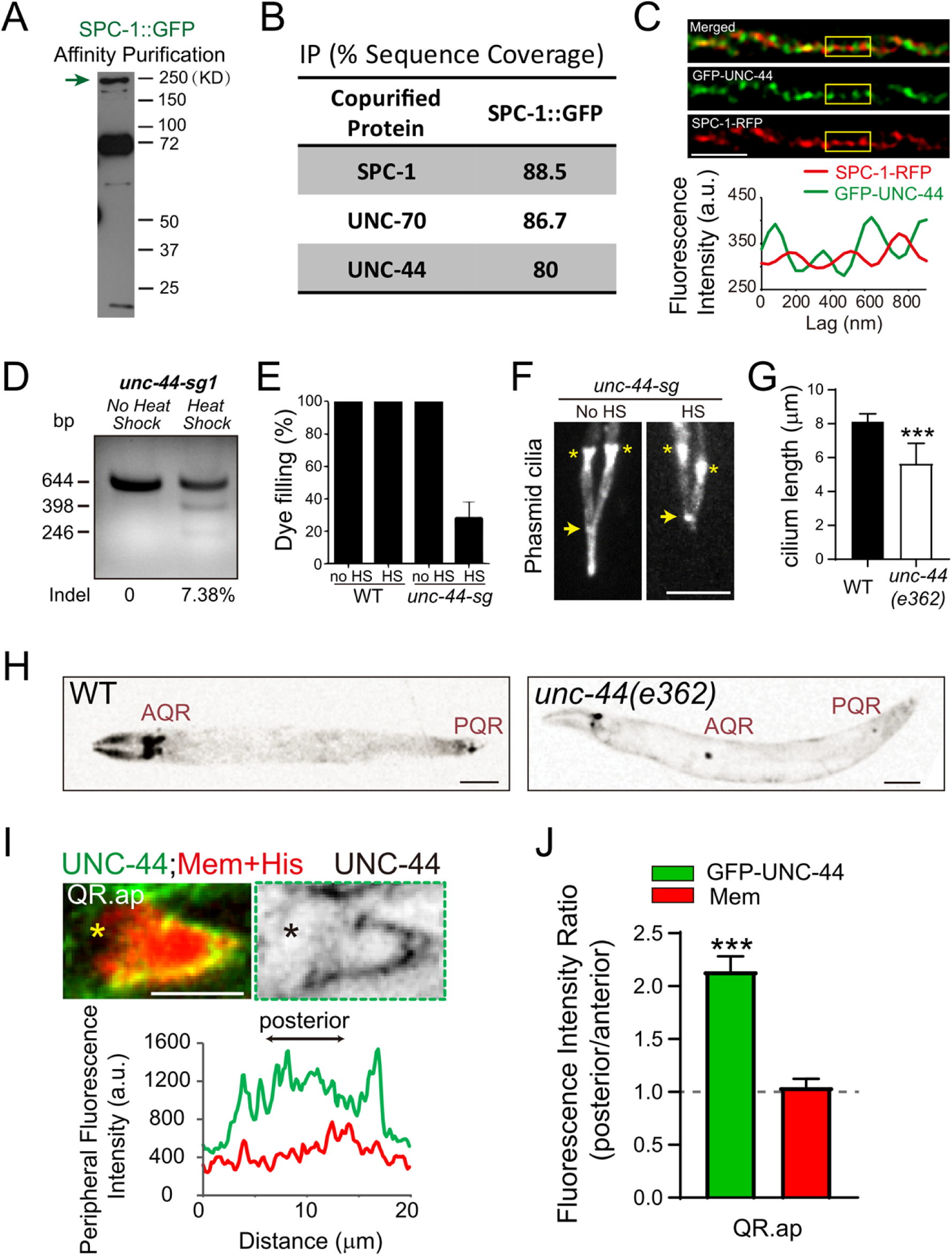
UNC-44 interacts with SPC-1 and regulates neuroblast development. **(A)** Western blot with anti-GFP antibody showing the immunoprecipitation of SPC-1::GFP (indicated in the gel with the green arrow) from *C. elegans* using GFP-TrapA beads. **(B)** Mass spectrometry analysis of the SPC-1::GFP affinity purifications. The percentage of sequence coverage was listed. **(C)** The representative image of GFP::UNC-44 (green) and SPC-1::RFP (red) in double knock-in animals (upper, merged; middle, GFP::UNC-44; and lower, SPC-1::RFP); bottom, the corresponding GFP::UNC-44 (green) and SPC-1::RFP (red) fluorescence intensity of the boxed region. Scale bar, 1 μm. **(D)** Representative gels of the T7EI assay for *unc-44* PCR products amplified from the genomic DNA of worms expressing P*hsp::Cas9* and P*U6::unc-44-sg1* with or without heat-shock treatment. **(E)** Dye-filling defects under a fluorescence stereoscope (*N* >100 animals) of WT animals or worms expressing P*hsp::Cas9* and P*U6::unc-44-sg1*, with or without heat-shock treatment, respectively. **(F)** Cilium morphology in conditional *unc-44* mutants, with or without heat-shock treatment, visualized by an intraflagellar transport protein IFT52/OSM-6::GFP. Arrows indicate the positions where the junctions between the middle and distal ciliary segments should be in WT animals. Asterisk, the ciliary base and transition zone. Scale bar, 5 μm. **(G)** Quantifications of the cilium length. **(H)** Fluorescence inverted images of the A/PQR position in WT and *unc-44(e362)* mutant animals. A/PQR neurons were visualized using P*gcy-32::mCherry*. The image is inverted so that high mCherry fluorescence intensity is black. The cell identities are denoted adjacent to the cells. Scale bar, 50 μm. **(I)** The representative images of GFP-tagged UNC-44 in QR.ap cell in knock-in animals, upper left showing GFP KI fluorescence (green) with mCherry-tagged plasma membrane and histone (red), upper right showing inverted fluorescence images of GFP KI fluorescence, and bottom showing the GFP (green) and mCherry (red) fluorescence distribution plot along the cell periphery of the corresponding imaging shown above. Asterisk, the plot start point. Scale bar, 5 μm. **(J)** Quantification of the GFP (green) tagged UNC-44 and mCherry (red) fluorescence intensity ratio between the posterior and the anterior membrane potions in the migrating QR.ap cells (N = 14 animals). The error bars indicate the standard error of the mean (SEM). Statistical significance is based on Student’s *t-*test, *** *P* < 0 .001.

Using the *unc-44(e362)* allele or conditional *unc-44* mutants generated by somatic CRISPR-Cas9, we showed that the loss of UNC-44 reduced Q cell migration in adult animals and disrupted the neurites in Q cell progenies as those in the spectrin mutant animals (Fig. 4H). By creating a UNC-44::GFP knock-in animal, we showed that UNC-44 was absent from the leading edge but accumulated in the lateral and the rear of migrating Q cells (Fig. 4I, and Movie S4), which is similar to spectrin distribution. We have previously reported the function of spectrin in ciliogenesis. In further support of the notion that UNC-44 acts with spectrin in the same processes, we found that the ciliary length and the capacity of the animal to interact with the environment (i.e., dye-filling) were defective in the loss of UNC-44 (Fig. 4E-G). Using super-resolution live imaging, we showed that UNC-44 displayed periodicity in the axon of motor neurons (Fig. 4C). Double fluorescence imaging analysis revealed that the fluorescence intensity peaks from UNC-44::GFP and SPC-1::mCherry showed intercalation in the axon (Fig. 4C), which is consistent with the current model on the organization of the spectrin-ankyrin skeleton (3, 8, 27). These results show that ankyrin functions together with spectrin in neuronal cell migration, the formation of the dendrite, and sensory cilia.

## Discussion

We provide evidence that the spectrin-ankyrin-based membrane skeleton is essential for neuronal cell migration and dendrite formation. This work reveals that the membrane skeleton is asymmetric and remodels during neural development: When neuroblasts migrate, spectrin accumulates in the lateral and the rear; when neuroblasts use the same leading-edge to grow dendrite, spectrin transits from the soma to the nascent dendrites. In comparison to the polarized and dynamic actin and microtubule cytoskeleton, asymmetry and remodeling are not well appreciated for the membrane skeleton. The previous studies on spectrin used “snapshot” imaging or fixed samples and described the membrane skeleton as an immobile and stable structure underneath the entire plasma membrane (3, 6, 8, 27). In addition to its rigidity, asymmetry and remodeling can be essential for the function of the spectrin-based membrane skeleton. A polarized membrane skeleton provides the locally distinct membrane mechanics in a living cell (e.g., cell migration), and the ability to remodel enables a developing cell to change its membrane mechanics at different developmental events (e.g., from cell migration to neurite outgrowth).

How can the spectrin-based membrane skeleton be asymmetric? Fluorescence microscopy of the cell from birth to migration uncovers that the membrane skeleton becomes asymmetrically assembled in the posterior (Fig. 2F). The inhibition of spectrin localization in the leading edge may result from some unknown chemical cues. Alternatively, the space beneath the plasma membrane of the leading edge is occupied by the Arp2/3-nucleated branching actin cytoskeleton (2), the dense network of which may serve as a physical barrier to block the assembly of the spectrin-based membrane skeleton. The similar physical inhibition may be applied out of the front where the membrane skeleton prevents the recruitment of the Arp2/3 and its activating protein machinery, blocking the polymerization of the branched actin network in the lateral and the rear.

Equally intriguing is how to remodel the assembled membrane skeleton. Despite immense knowledge on the depolymerization of actin filaments and microtubule, little is known regarding the disassembly of the membrane skeleton. The extracellular signal-regulated kinase signaling caused calpain-dependent degradation of the membrane-associated periodic structure, which suggests that the kinase activity may be involved (9). A protein kinase within the lateral and rear of the migrating cell may phosphorylate spectrin and disassemble the membrane skeleton at the end of cell migration. Our early study identified that the *C. elegans* Hippo kinases localize outside of the front and phosphorylate a small GTPase RhoG/MIG-2, inhibiting its activity from activating Arp2/3 (18). Besides, a MAPK family kinase MAP4K4 in the rear phosphorylates the actin-membrane linker ERM during mammalian cell migration (30). Such kinases may be involved in remodelling the membrane skeleton when the cell completes its movement. Alternatively, other kinases or other types of protein posttranslational modifications may regulate the disassembly of the membrane skeleton.

The spectrin-based membrane skeleton and the branched actin network appear to be mutually exclusive. The branched actin network assembles in the leading edge where spectrin is absent, and actin is ectopically assembled in the posterior of *spc-1* deficient cells (Fig. 1F-G and Fig. 2C-G). The Arp2/3-stimulated actin polymerization, Coronin-based F-actin debranching, and ADF/Cofilin-mediated F-actin severing empower a highly dynamic and protrusive membrane in the leading edge (2). The rigidity of the spectrin-based membrane skeleton offers stable membrane mechanics that maintain cell morphology of the migrating cells and the newly formed dendrite. The protrusive and rigid membrane domains in the same developing cell are thus well organized by the coordinated action of Arp2/3 and spectrin. Such a negative correlation between the branched actin assembly and the spectrin-based membrane mechanics provides insights into a long-standing question of how a migrating cell forms only one leading edge. A prevailing model posits that the plasma membrane tension constrains the spread of the leading edge and prevents the formation of new fronts. For example, migrating neutrophils use plasma membrane tension as a physical inhibitor to impede actin assembly outside of the existing front (31), but the molecular basis of physical tension is unclear. The spectrin-based membrane skeleton may play a previously unrecognized yet potentially widespread role in membrane tension during different types of cell migration.

The findings that the hereditary elliptocytosis-associated α-spectrin mutation and the spinocerebellar ataxia type 5-associated β-III–spectrin deletion are defective in neuronal cell migration and neurite formation support the notion that the spectrin-based membrane mechanics plays multiple roles in neural development, providing additional insights into SCA5 pathogenesis. Our results will motivate future investigations to examine whether neuronal migration or axon and dendrite formation are abnormal in Purkinje cells and granule cell progenitors with hereditary elliptocytosis or spinocerebellar ataxia. The manipulation of mechanobiology in the nervous system may merit exploration as a potential therapeutic strategy for some types of neurodegenerative disorders.

## Materials and Methods

### *C. elegans* strains and genetics

*C. elegans* strains were raised on NGM plates seeded with *E. coli* strain OP50 at 20°C. Tables S1-S3 summarizes the primers, plasmids, and strains used in this study.

### Molecular biology

CRISPR-Cas9 targets were inserted to the pDD162 vector (Addgene #47549) by linearizing this vector with primers listed in Table S1. The resulting PCR products containing 15 bp overlapped double-strand DNA ends were treated with DpnI digestion overnight and transformed into *E. coli*. The linearized PCR products were cyclized to generate plasmids by spontaneous recombination in bacteria. For fluorescence tag knock-in, homology recombination (HR) templates were constructed by cloning the 2 kb 5’ and 3’ homology arms into pPD95.77 plasmids using In-Fusion Advantage PCR cloning kit (Clontech, cat. no. 639621). We used the CRISPR design tool (http://crispr.mit.edu) to select the target sequence. GFP, GFP_11×7,_ and RFP tags were added to the C-terminus of SPC-1, whereas UNC-70 and UNC-44 were tagged with an N-terminal GFP. We generated SPC-1-L268P and UNC-70-ΔH590-L598 mutation knock-ins in the SPC-1::GFP and GFP::UNC-70 knock-in background by the similar cloning strategies. Conditional knockout strains were generated and determined as previously described (Shen et al., 2014).

### Microinjection and transgenesis

For SPC-1 split GFP knock-in generation, P*egl-17*::GFP_1-10_ constructs were generated by cloning *egl-17* promotor and GFP_1-10_ sequences into pPD95.77 plasmids using In-Fusion Advantage PCR cloning kit (Clontech, cat. no. 639621). The plasmid was injected into SPC-1::GFP_11×7_ knock-in animals. Transgenic strains were generated by microinjecting ∼50 ng/μl DNA with co-injection markers into *C. elegans* germlines.

### Mass spectrometry analysis

Unsynchronized SPC-1::GFP knock-in strains raised on 80∼100 90-mm NGM plates were collected and washed for three times with M9 buffer. Lysates were made from 1∼2 mL packed worms in lysis buffer (25 mM Tris-HCl, pH 7.4, 150 mM NaCl, 1% NP-40, 10% glycerol, 1x cocktail of protease inhibitors from Roche (Complete, EDTA free), 40 mM NaF, 5 mM Na3VO4) and 3∼4 mL of 0.5-mm diameter glass beads using FastPrep-24 (MP Biomedicals). Proteins were immunoprecipitated with GFP-Trap A beads (Chromoteck) and eluted with 300 mL 0.1 M glycine-HCl, pH 2.5 into 15 mL 1.5 M Tris-HCl pH 8.8, followed by precipitation with 100 mL trichloroacetic acid and redissolved in 60 mL 8 M urea, 100 mM Tris-HCl, pH 8.5. Samples were treated with 5 mM TCEP for reduction, 10 mM iodoacetamide for alkylation, and then diluted fourfold with 100 mM Tris-HCl 8.5. The proteins were digested with 0.2 mg trypsin at 37°C overnight after the addition of 1 mM CaCl2, and 20 mM methylamine. The resultant peptides were desalted with Zip Tip pipette tips (Merck Millipore). The tandem mass spectrometry spectra were searched against the *C. elegans* proteome database using Proteome Discoverer (version PD1.4; Thermo Fisher Scientific). The protein components in the list in Figure 4 were reproducibly identified from three biological repeats.

### Dye-filling assay

Young-adult worms were randomly collected into 200 μl M9 solution and mixed with equal volume dyes (DiI, 1,1’-dioctadecyl-3,3,3’,3’,-tetramethylindo-carbocyanine perchlorate; Sigma-Aldrich, St. Louis, MO, USA) at working concentration (20 μg/ml), followed by incubation at room temperature in the dark for 30 min. Worms were transferred to seeded NGM plates and examined for dye uptake one hour later using a fluorescence stereoscope. At least 100 worms of each strain were examined in two independent assays.

### Live-cell imaging

Q cell migration imaging was performed using the previously described protocol (32). L1 worms were anesthetized with 0.1 mmol/L levamisole in M9 buffer and mounted on 3% agarose pads at 20°C, then imaged on an Axio Observer Z1 microscope (Carl Zeiss) equipped with a 100×, 1.49 numerical aperture (NA) objective, an electron-multiplying (EM) charge-coupled device (CCD) camera (Andor iXon+ DU-897D-C00-#BV-500), and the 488 nm and 561 nm lines of a Sapphire CW CDRH USB Laser System attached to a spinning disk confocal scan head (Yokogawa CSU-X1 Spinning Disk Unit). Time-lapse images were acquired by μManager (https://www.micro-manager.org) at an exposure time of 200 msec. Images of spectrin and UNC-44 periodicity were collected on a Nikon (Tokyo, Japan) A1R laser-scanning confocal microscope with a CFI Plan Apo 100× oil immersion objective (numerical aperture 1.45) and 488-nm lasers with 150 nm X-Y resolution.

### Quantifications and statistical analysis

ImageJ software was used to circumscribe the fluorescence field and to measure the fluorescence intensity. In all the intensity quantifications, the background was subtracted. When the F-actin marker GFP::moesinABD or knock-in fluorescence protein signal intensity was measured along the cortex, “leading” refers to the circumference of the lamellipodium (the wide-spread area) and “rear” refers to the circumference around the rest of cell body, and the intensity ratio between the two was calculated using the corresponding mCherry fluorescence intensity as a control. The migration distances were measured as the cell body movement (in μm per hour) along the anterior-posterior body axis. The migration angle was measured between the protrusion and the anterior-posterior body axis, the anterior was defined as degree 0, and the posterior was defined as degree 180. Quantification was represented by the mean value ± standard deviation for each group. N represents the number of animals used for the corresponding quantification, and the one-sample or two-tailed Student’s *t-*test or χ^2^ analysis were employed to determine the statistical differences as indicated in figure legends.

## Acknowledgments

This work was supported by the National Key R&D Program of China (2017YFA0102900), the National Natural Science Foundation of China (grants 31730052, 31525015 and 31561130153, 31671444, 31871352), Beijing Municipal Natural Science Foundation (5172015).

## Supplemental Movie Legends

**Figure S1.**
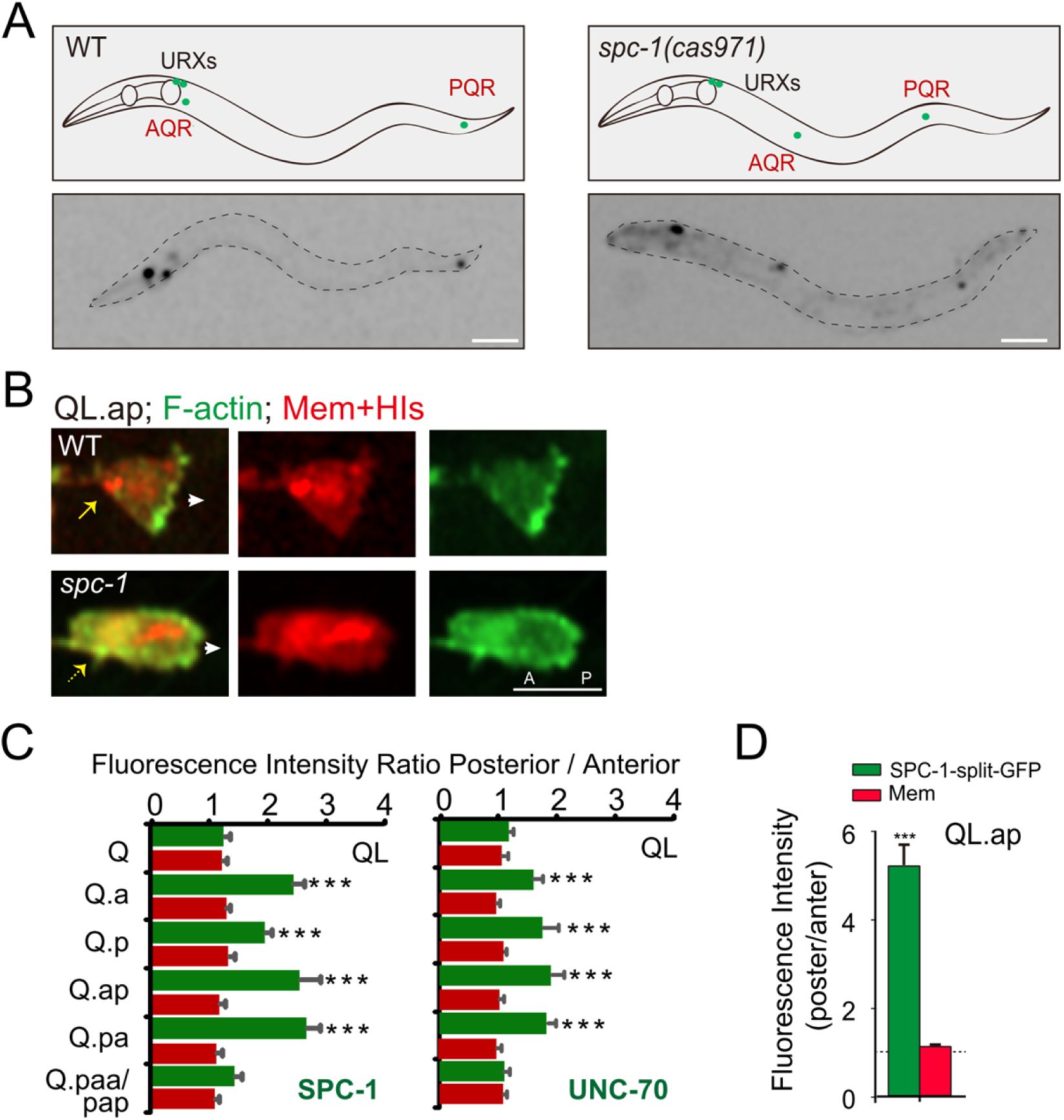
**(A)** Schematics (upper) and fluorescence inverted images of the A/PQR position in WT and *spc-1(cas971)* mutant animals. A/PQR neurons were visualized using P*gcy-32::mCherry*. The image is inverted so that high mCherry fluorescence intensity is black. The cell identities are denoted adjacent to the cells. Dotted blue lines show the periphery of *C. elegans*. Scale bar, 50 μm. **(B)** Fluorescence images of GFP-tagged F-actin (green) with mCherry (red) labeled plasma membrane and histone in QL.ap cells in WT or *spc-1(cas971)* animals. Yellow arrows show the rear of migrating cells; White arrows indicate the direction of migration. AP, anterior, and posterior. Scale bar, 5 μm. **(C)** Quantification of the GFP (green) of SPC-1 (left) or UNC-70 (right), and mCherry (red) fluorescence intensity ratio between the posterior and the anterior plasma membrane portions in the QL cell lineages (N = 10-20 animals). Anterior and posterior were divided by a dashed line in Fig. 2A. **(D)** Quantification of the fluorescence intensity ratio of split-7xGFP-tagged SPC-1 (green), and mCherry-membrane (red) between the posterior and the anterior of the migrating QL.ap cells (N = 15 animals). The error bars indicate the standard error of the mean (SEM). Statistical significance is based on Student’s *t-*test, *** *P* < 0 .001.

**Table S1.**
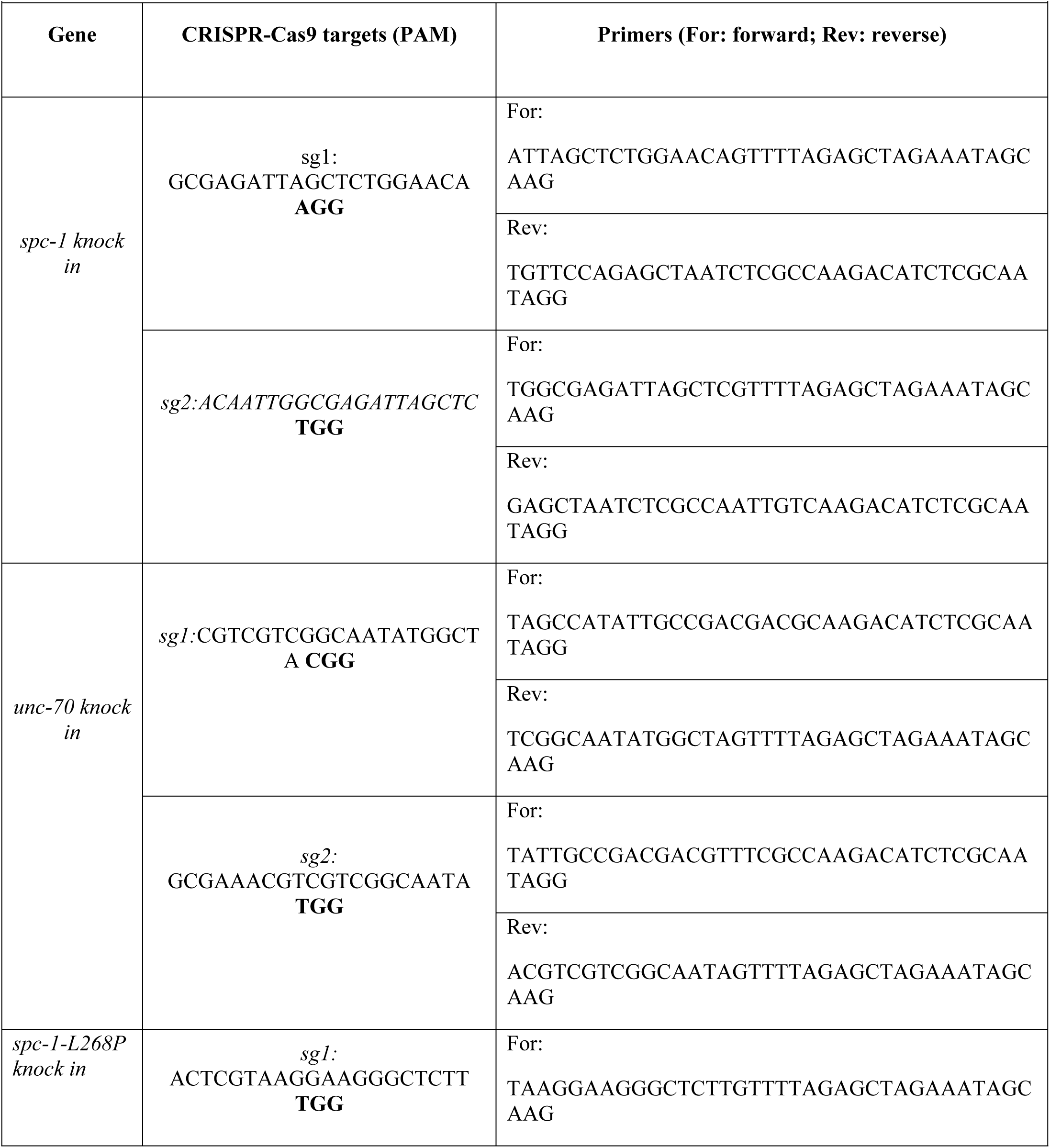
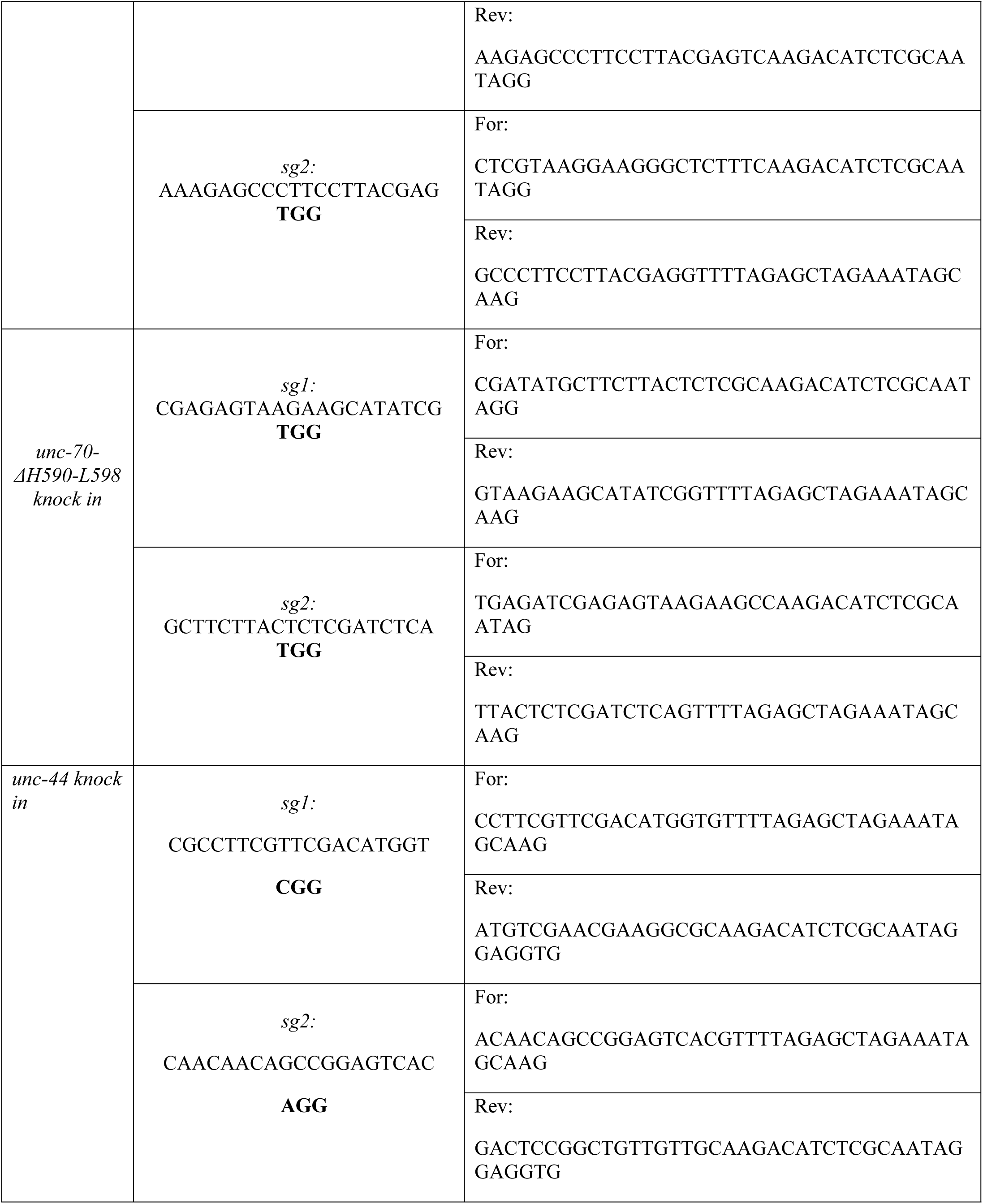
Targets of CRISPR and primers for molecular analysis.

**Table S2.**
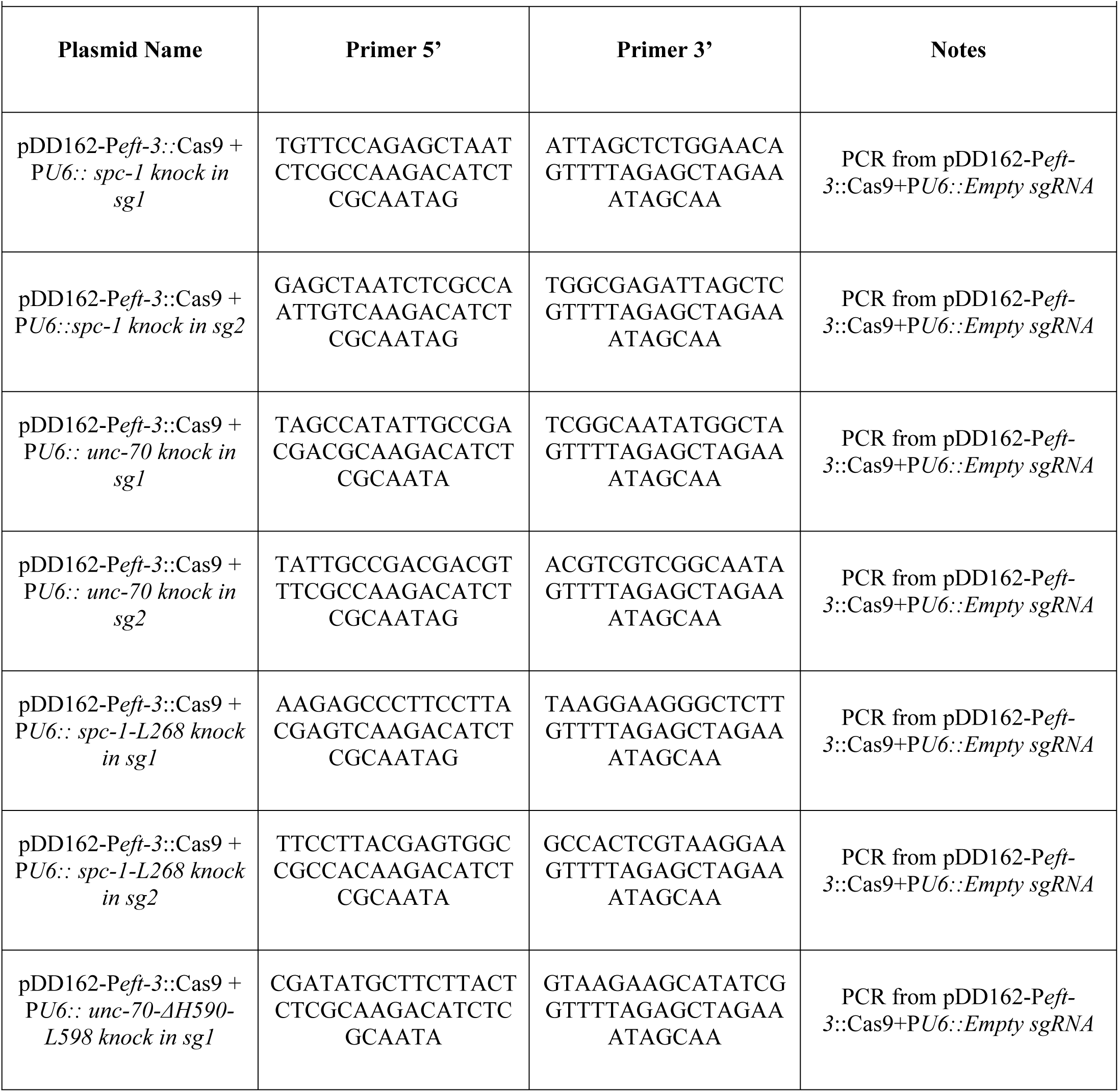
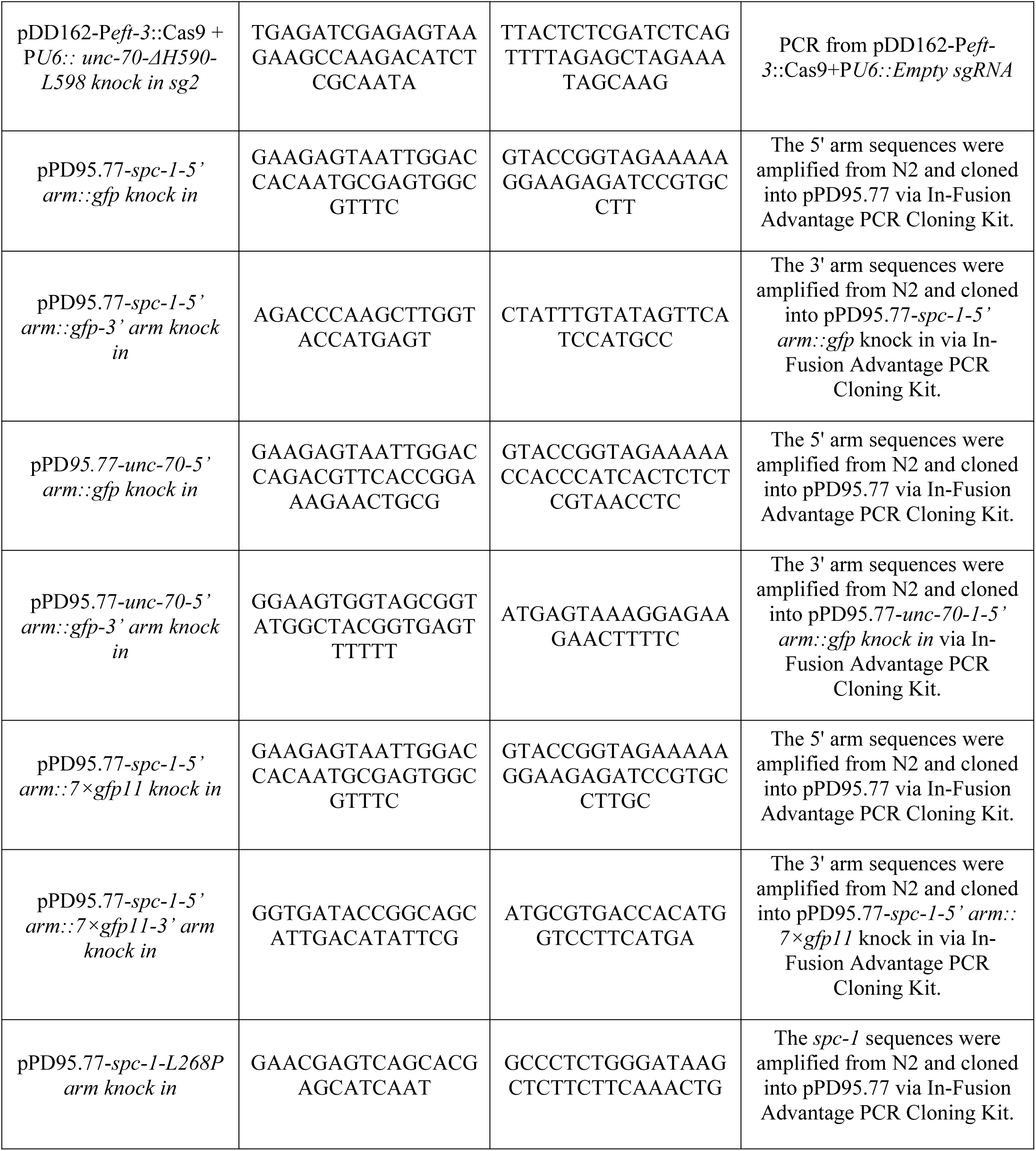
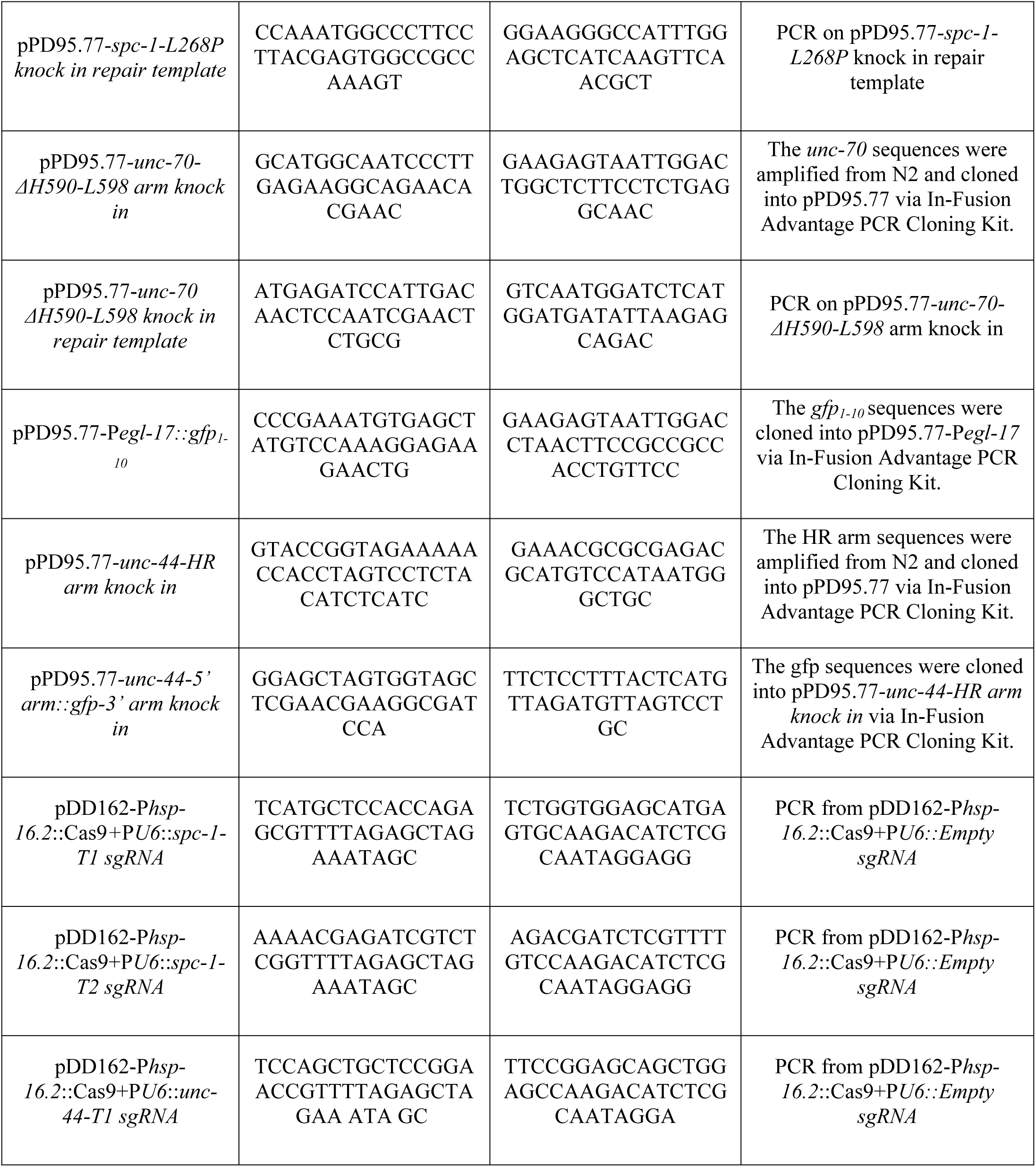
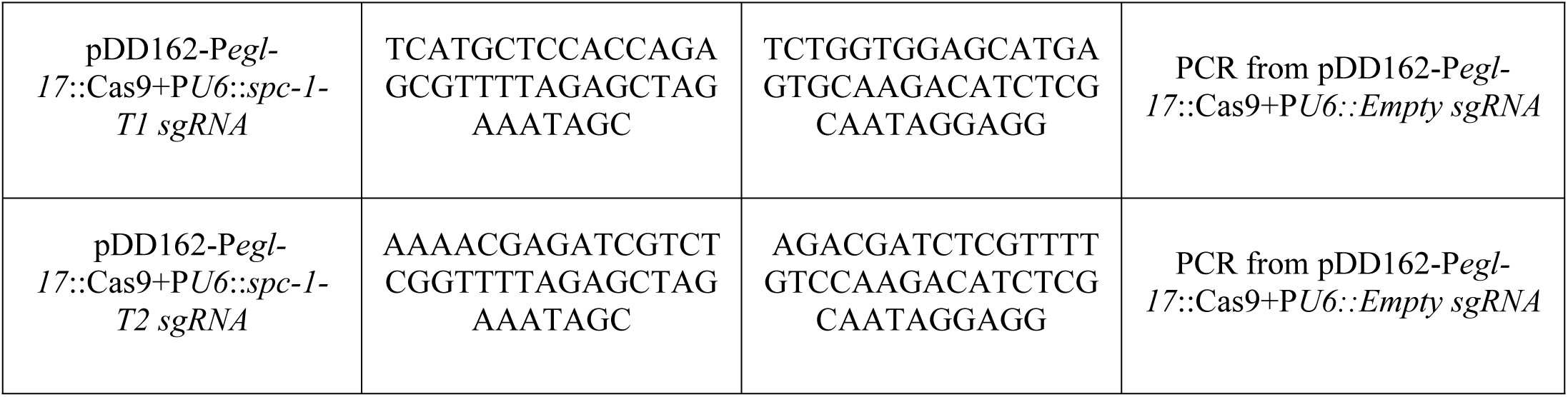
Primers and plasmids used for plasmid cloning in this study.

**Table S3.** *C. elegans* strains used in this study.

| Strain name | Genotype | Method | Resource |
| --- | --- | --- | --- |
| N2 | Wild type | - | CGC |
| GOU2936 | *cas815[spc-1::gfp knock in]* | Microinjection | This study |
| GOU3238 | *cas971[spc-1-L260P knock in]* | Microinjection | This study |
| GOU3519 | *cas1047[spc-1::7×gfp11 knock in]* | Microinjection | This study |
| GOU3617 | *cas1047[spc-1::7×gfp11 knock in]; casEx5751[Pegl-17::gfp1-10,pRF4(+)]* | Genetic cross | This study |
| GOU3103 | *cas962[gfp::unc-70 knock in]* | Microinjection | This study |
| GOU3237 | *cas983[unc-70-ΔH590-L598 knock in]* | Microinjection | This study |
| GOU2039 | *cas963[gfp::unc-44 knock in]* | Microinjection | This study |
| GOU3659 | *cas963[gfp::unc-44 knock in]; cas961[spc-1::rfp knock in]* | Genetic cross | This study |
| GOU174 | *casIs35[Pgcy-32::mCherry, unc-76(+)] ;*<br>*zdIs5[Pmec-4::gfp, lin-15(+)]* | Genetic cross | This study |
| GOU3648 | *casEx5752[Pegl-17::Cas9+PU6::spc-1-sg2,pRF4 ];*<br>*casIs35[Pgcy-32::mCherry, unc-76(+)];*<br>*zdIs5[Pmec-4::gfp, lin-15(+)]* | Microinjection | This study |
| GOU3649 | *casEx5753[Phsp16.2::Cas9+PU6::spc-1-sg1,pRF4 ];*<br>*casIs35[Pgcy-32::mCherry, unc-76(+)];*<br>*zdIs5[Pmec-4::gfp, lin-15(+)]* | Microinjection | This study |
| GOU3651 | *casEx5754[Phsp16.2::Cas9+PU6::spc-1-sg2,pRF4 ];*<br>*casIs35[Pgcy-32::mCherry, unc-76(+)];*<br>*zdIs5[Pmec-4::gfp, lin-15(+)]* | Microinjection | This study |
| GOU3072 | *cas971[spc-1-L260P knock in]; casIs35[Pgcy-32::mCherry,*<br>*unc-76(+)];* *zdIs5[Pmec-4::gfp, lin-15(+)]* | Genetic cross | This study |
| GOU3079 | *unc-44(e362); casIs35[Pgcy-32::mCherry, unc-76(+)];*<br>*zdIs5[Pmec-4::gfp, lin-15(+)]* | Genetic cross | This study |
| GOU4190 | *unc-44(e362); casEx5752[Pegl-17::Cas9+PU6::spc-1-*<br>*sg2,pRF4 ];* *casIs35[Pgcy-32::mCherry, unc-76(+)];*<br>*zdIs5[Pmec-4::gfp, lin-15(+)]* | Genetic cross | This study |
| GOU2937 | *cas815[spc-1::gfp knock in]; casIs165[Pegl-17:: myri-*<br>*mCherry, Pegl-17::mCherry-TEV-S::his-24, unc-76(+)]* | Genetic cross | This study |
| GOU4191 | *cas962[gfp::unc-70 knock in]; casIs165[Pegl-17:: myri-*<br>*mCherry, Pegl-17::mCherry-TEV-S::his-24, unc-76(+)]* | Genetic cross | This study |
| GOU3065 | *cas963[gfp::unc-44 knock in]; casIs165[Pegl-17:: myri-*<br>*mCherry, Pegl-17::mCherry-TEV-S::his-24, unc-76(+)]* | Genetic cross | This study |
| GOU1544 | *casIs165[Pegl-17:: myri-mCherry, Pegl-17::mCherry-TEV-*<br>*S::his-24, unc-76(+)];* *casIs555[Pegl-17::gfp::moesinABD]* | Genetic cross | This study |
| GOU3676 | *cas971[spc-1-L260P knock in]; casIs165[Pegl-17:: myri-mCherry, Pegl-17::mCherry-TEV-S::his-24, unc-76(+)] II; casIs555[Pegl-17::gfp::moesinABD]* | Genetic cross | This study |
| GOU1409 | *unc-44(e362); mnIs17[osm-6::gfp]* | Genetic cross | This study |
| GOU2035 | *casEx2000[Phsp-16.2::Cas9+PU6::unc-44-T3 sgRNA; Podr-1::dsRed; ]; mnIs17[osm-6::gfp]* | Genetic cross | This study |

**Movie S1 is related to Figure 2. Dynamic distribution of SPC-1::-7xGFP during QR**.**ap cell migration**

Fluorescence time-lapse movies of QR.ap cell migration with 7xGFP-tagged SPC-1 and mCherry-tagged plasma membrane in a WT animal. Frames were taken every 60 seconds. The display rate is 7 frames per second.

**Movie S2 is related to Figure 3. The neuritogenesis of QL**.**ap in WT**

Fluorescence time-lapse movies of GFP-tagged F-actin and mCherry-tagged plasma membrane during QL.ap neuritogenesis, showing the transformation of the leading edge into the growth cone. Frames were taken every 60 seconds. The display rate is 7 frames per second.

**Movie S3 is related to Figure 3. The neuritogenesis of QL**.**ap in *spc-1(cas971)* mutant animal**

Fluorescence time-lapse movies of GFP-tagged F-actin and mCherry-tagged plasma membrane during QL.ap neuritogenesis in *spc-1(cas971)* mutants, showing the transformation of the leading edge into the branched growth cone. Frames were taken every 60 seconds. The display rate is 7 frames per second.

**Movie S4 is related to Figure 4. Dynamic distribution of GFP::UNC-44 during QR**.**a and QR**.**ap cell migration in the knock-in animals**

Fluorescence time-lapse movies of QR.ap cell migration with GFP-tagged UNC-44 and mCherry-tagged plasma membrane in a knock-in animal. Frames were taken every 60 seconds. The display rate is 7 frames per second.

